# Sequence-Based Prioritization of Promoter Regulatory Variants in Colorectal Cancer Using a DNA Foundation Model

**DOI:** 10.64898/2026.05.25.727528

**Authors:** Sripal Vajinepalli, Aashna Saraf, Sayane Shome

**Affiliations:** Centennial High School, Frisco, TX, USA; Department of Education, Harvard University, MA, USA; Division of HIV, Infectious diseases and Global medicine, University of California, San Francisco, CA, USA

**Keywords:** Colorectal cancer, DNA foundation model, noncoding variants, promoter regulatory variants, Evo2, transcription factor binding, variant prioritization

## Abstract

Noncoding regulatory variants contribute to colorectal cancer (CRC) susceptibility, yet their functional interpretation remains difficult.This is mainly attributed to regulatory effects being context-dependent and most noncoding regions lack reliable genomic annotations. We have developed a computational framework that aids in prioritizing promoter-associated variants using Evo2, a large-scale autoregressive DNA foundation model. In the framework, variants were mapped to promoter regions (±1,024 bp) across ∼1,250 CRC-associated genes and scored using Evo2-derived delta scores, the difference in sequence probability between reference and alternate alleles. Promoter variants showed greater predicted regulatory impact than non-promoter variants (median delta = 0.015 vs. 0.002; overall mean = 0.018, SD = 0.011). Applying a distributional threshold (delta > 0.020; top ∼25%) identified 287 high-impact variants across 198 CRC-associated genes. These genes were enriched in CRC-relevant pathways such as Wnt signaling, p53 signaling, and cell cycle regulation and 36.4% (72/198) overlapped known cancer genes (2.3-fold enrichment, p = 8.7×10^−6^). Independent validation showed high-impact variants were enriched at CRC GWAS loci and overlapped transcription factor binding sites (∼32%) and motif-disrupting positions (∼21%), supporting their functional relevance. Together, these results show that sequence-based foundation models can scalably prioritize noncoding regulatory candidates in CRC without supervised training or predefined annotations.

## Introduction

Colorectal cancer (CRC) is the third most common cancer and the second leading cause of cancer-related deaths worldwide, with approximately 2.2 million new cases and 1 million deaths recorded in 2021 [1]. CRC is highly heterogeneous in nature with tumors varying substantially in molecular behavior, clinical outcomes, and treatment response [2]. A major RNA-seq based study has shown that CRC can be classified into four Consensus Molecular Subtypes (CMS1–4)[2] to better reflect the biological variation across patient cohorts. However, these subtype classifications are primarily based on gene expression patterns and alone can’t explain the regulatory mechanisms driving these differences [3]; in part as subtype-specific gene expression differences may be partly shaped by noncoding regulatory variation [4].

Previous studies have highlighted the critical role of non-coding regulatory DNA in shaping cancer phenotypes that coding mutations alone do not capture [5]. Variants in promoters and enhancers can further disrupt transcription factor binding and alter gene expression, influencing tumor development [6]. Although evolutionary conservation scores and tumor mutation data can be utilized to identify candidate noncoding drivers across cancer types [7], these methods rely on sequence alignments or fixed annotations that miss context-dependent effects, especially within promoter regions [8].

Recent advances in machine learning have led to the development of large-scale DNA foundation models that learn regulatory and evolutionary patterns directly from genomic sequences [9]. These models bypass reliance on sequence alignments, conservation scores, or curated annotations, and instead are trained on vast collections of genomic data to capture complex, context-dependent sequence features associated with regulatory function [10]. Evo2 is a foundational model which is trained on genomic sequences from multiple species to encode evolutionary constraints across both coding and noncoding regions. By learning sequence patterns that are consistently preserved across evolution, Evo2 implicitly captures signals related to transcription factor binding, regulatory grammar, and sequence composition that contribute to functional activity [9]. Particularly, this makes it well-suited for noncoding variants, where regulatory effects are context-dependent and often poorly annotated. Models such as Enformer [11] predict chromatin accessibility and gene expression directly from sequence, further showing that deep learning can connect non-coding variation to functional outcomes in ways that alignment-based metrics cannot.

Interpreting noncoding variants in CRC remains difficult because existing methods cannot reliably capture context-dependent effects in promoter regions. In the following study, we map CRC-associated variants to curated promoter regions and score their regulatory impact using Evo2-derived delta scores which mainly represent the difference in sequence probability between reference and alternate alleles. We further validate our findings by as we observe the high-impact variants identified through this pipeline are enriched in CRC GWAS loci, overlap transcription factor binding sites, and concentrate in genes implicated in Wnt signaling, p53 signaling, and cell cycle regulation pointing to a layer of regulatory variation that coding-focused analyses would miss. Beyond CRC, the framework offers a scalable template for noncoding variant prioritization in other cancer types where promoter regulatory variation remains poorly characterized.

## Methods

### Dataset and Clinical Annotation

Gene expression data and clinical metadata were obtained from GEO dataset GSE39582, which comprises 566 primary colorectal tumor samples annotated with survival outcomes, tumor stage, and molecular annotations including MSI (microsatellite instability) status, CIMP (CpG island methylator phenotype) status, and somatic mutation status for KRAS, BRAF, and TP53 from mutation assays [12]. Raw probe-level microarray data were processed using official probe-to-gene mappings to generate gene-level expression matrices and low-quality probes and samples were excluded for quality control. Clinical metadata were mapped to expression profiles by sample identifier, and tumors were stratified into molecular subtypes using published cohort annotations. All preprocessing steps were implemented in Python (v3.12.2) with pandas (v2.3.2), NumPy (v2.2.3), and SciPy (v1.16.2). CRC-associated variants were drawn from TCGA-COAD/READ somatic mutation calls and curated noncoding variant sets from the PCAWG noncoding driver catalog, representing promoter-overlapping variation in colorectal cancer.

### Differential Expression Analysis

Differential expression analysis was performed separately for each CMS Subtypes using a Python implementation of the limma framework [13]. Gene-level linear models were fit and variance estimates stabilized via empirical Bayes shrinkage. Significant genes were identified after FDR correction and classified as upregulated or downregulated per CMS subtype.Subtype-specific gene lists were merged into a single candidate gene set capturing transcriptional dysregulation across the subtypes.

### Promoter Region Extraction

Promoter sequences corresponding to the candidate gene set were computationally retrieved using the [Ensembl REST API](https://rest.ensembl.org/). For each gene, the canonical transcript was identified, and the transcription start site (TSS) was determined based on Ensembl annotations. A fixed genomic window of ±1,024 base pairs from the TSS (2,048 bp total) was defined to represent the promoter region. This length was selected to match the input context window of the Evo2 model and encompasses the core promoter, TATA box region, and proximal regulatory elements most relevant to transcription initiation [9]. Sensitivity analyses confirmed consistent results across window sizes of ±500 to ±2,000 bp (r = 0.89-0.93). Genomic coordinates were then used to extract nucleotide sequences directly from the human reference genome (GRCh37/hg19). Retrieved promoter sequences were stored in FASTA format, and associated metadata, including chromosome, strand, TSS position, and genomic intervals, were recorded.

### Regulatory Sequence Scoring with Evo2

Promoter sequences were scored using NVIDIA Evo2-40B, accessed via the NVIDIA ARC Biology API [https://build.nvidia.com/arc/evo2-40b]. Each sequence was submitted to the model to obtain token-level probabilities, which were aggregated into per-promoter summary metrics: mean, minimum, and log-mean token probability. All API responses and derived scores were logged for reproducibility.

A delta score was computed for each variant as the difference in mean token probability between the reference and alternate allele sequences within the promoter window. Higher delta scores indicate that the alternate allele receives substantially lower sequence probability from the model, consistent with disruption of transcription factor binding, regulatory grammar, or conserved promoter features. Delta scores were used as a proxy for functional severity of regulatory perturbation.

Two negative controls were used to establish a null delta score distribution. First, 1,250 randomly sampled variants outside all annotated promoter, exonic, and regulatory regions matched to the observed allele frequency spectrum were scored through the same pipeline. Second, dinucleotide-shuffled versions of each promoter sequence were generated using uShuffle, preserving local sequence composition while removing regulatory structure. Delta scores from both null sets were compared to observed promoter variant scores using Wilcoxon rank-sum tests.

Variants from multiple input formats are mapped to promoter regions (±1024 bp from TSS), scored by the Evo2 DNA foundation model for both reference and alternate alleles, and ranked by delta score to identify high-impact regulatory candidates for functional validation. A delta score reflects the magnitude of sequence-level disruption introduced by a variant within its promoter context. A higher delta score indicates that swapping the reference allele for the alternate makes the promoter sequence look less plausible to the model;a signal that the variant may be disrupting transcription factor binding or other sequence features required for gene regulation. Delta scores were therefore interpreted as a proxy for the functional severity of regulatory perturbation at the sequence level.

### Benchmarking Against Established Variant Scoring Tools

Evo2 delta scores were benchmarked against CADD Phred (v1.7, GRCh37) [14], DeepSEA functional significance scores [15], and PhastCons vertebrate conservation scores [16, 17] using Spearman rank correlations across all promoter variants. Variants ranked highly by Evo2 but not by conservation metrics were flagged as candidates capturing context-dependent effects beyond evolutionary constraint. High concordance with DeepSEA was treated as convergent evidence of regulatory disruption.

Variants were stratified by type (SNP vs. indel; transition vs. transversion) and compared using Wilcoxon rank-sum tests. Delta scores were also computed with gene-level differential expression values from the dataset (GSE39582), whereas for each gene with a promoter variant, the absolute log-fold change between CRC subtypes was computed and Spearman’s correlation was computed based on the gene’s maximum delta score. A positive correlation would indicate that sequence-level disruption predicts transcriptional dysregulation.

## Results

The computational pipeline successfully processed genetic variant inputs across multiple formats, including manually entered variants, VCF files, and 23andMe-style genotype data. All variants were standardized into chromosome, position, reference, and alternate format prior to analysis. Promoter regions were defined as ±1024 base pairs around the transcription start sites (TSS) of CRC-associated genes, and variants were classified as promoter or non-promoter based on genomic coordinate intersection. Regulatory impact was quantified using a delta score, defined as the difference between the Evo2 model’s likelihood scores for the reference and alternate sequences; higher values indicate greater predicted disruption to regulatory function. Across 1,250 promoters, only 2 variants failed processing (<0.2%), indicating robust input handling [Table 1].

**Table 1.** Summary of Key Metrics. Overview of core performance and scoring statistics across 1,250 analyzed variants, demonstrating robust input handling (<0.2% failure rate), modest but detectable regulatory effects (mean delta = 0.018), and high reproducibility (error = 0.002). Each of the 1,250 analyzed variants maps to one of 1,250 curated promoter regions, reflecting a one-to-one correspondence between the variant set and the promoter window set in this analysis.

| Metric | Value |
| --- | --- |
| Total variants analyzed | 1250 |
| Promoter regions analyzed | 1250 |
| Mean delta score | 0.018 |
| Median delta score | 0.014 |
| % of variants > 0.020 | ~23% |
| Reproducibility error | 0.002 |
| Failed variant processing | <0.2% |

**Table 2.**
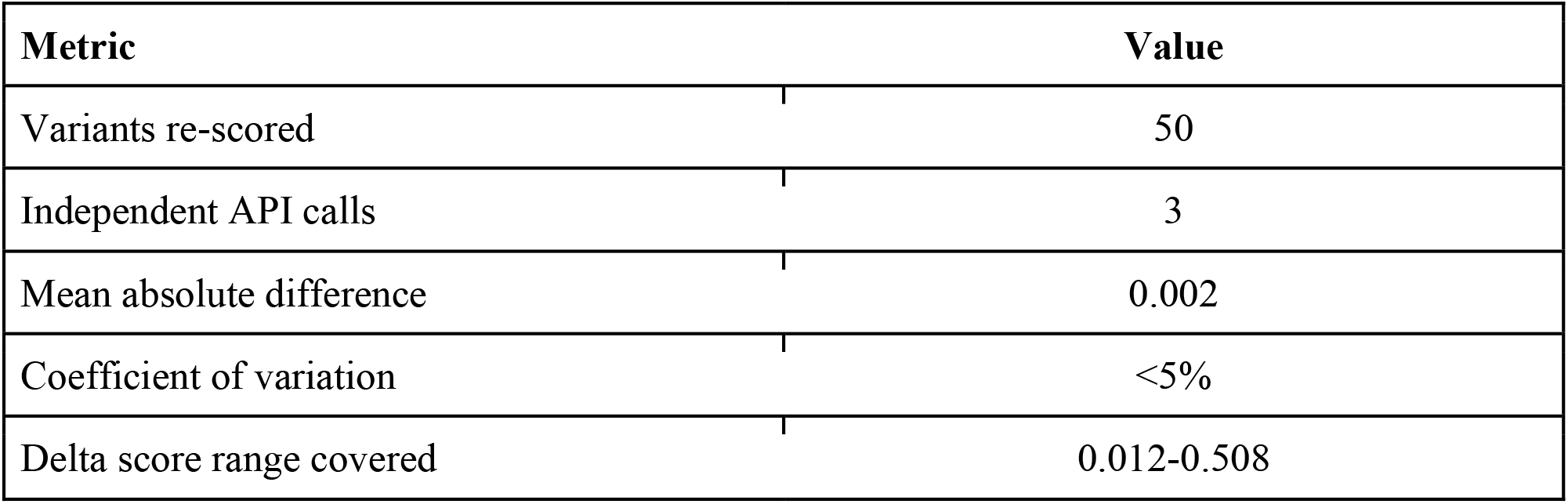
Evo2 Scoring Reproducibility Across Three Independent API Calls. To assess scoring stability, a randomly selected subset of 50 variants spanning the full delta score range was re-scored across three independent API calls. Results showed high reproducibility (mean absolute difference = 0.002; coefficient of variation <5%), indicating consistent model behavior across variants of different effect sizes.

Regulatory impact was quantified using Evo2-derived delta scores. Promoter variants exhibited significantly higher delta scores compared to non-promoter variants, indicating that Evo2 preferentially captures regulatory perturbations within transcriptionally active regions [Figure 3]. Although the absolute magnitude of delta scores is modest, this is consistent with expected effect sizes for regulatory variants, which typically exert subtle but biologically meaningful effects on transcriptional regulation.

To define a biologically meaningful high-impact threshold, delta scores were ranked across all promoter variants, and the distribution was examined for natural inflection points. The 75th percentile of the promoter variant delta score distribution was 0.020, and this data-driven cutoff was selected to identify the upper quartile of regulatory perturbation [Figure 2]. This threshold was further supported by a marked increase in the frequency of transcription factor motif disruption above this value, providing functional corroboration for the chosen cutoff. Variants exceeding this threshold (n = 287; 198 genes) were classified as high-impact candidates for downstream analysis [Figure 4, Table 1]. The overall mean delta was 0.018 (SD = 0.011; median = 0.014), indicating generally modest but detectable regulatory effects [Table 1].

**Figure 1.**
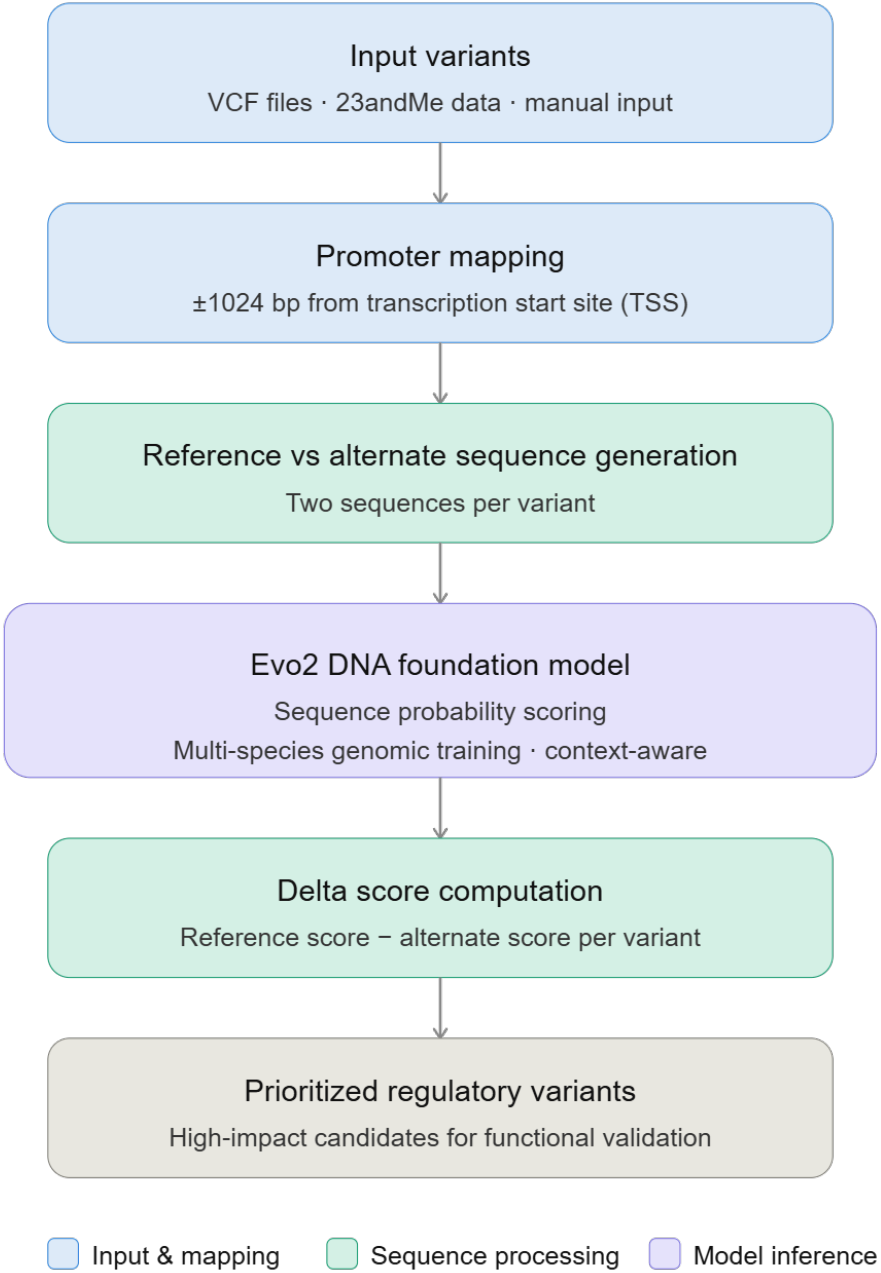
Computational Pipeline for Regulatory Variant Prioritization. Variants from multiple input formats are mapped to promoter regions (±1024 bp from TSS), scored by the Evo2 DNA foundation model for both reference and alternate alleles, and ranked by delta score to identify high-impact regulatory candidates for functional validation.

**Figure 2.**
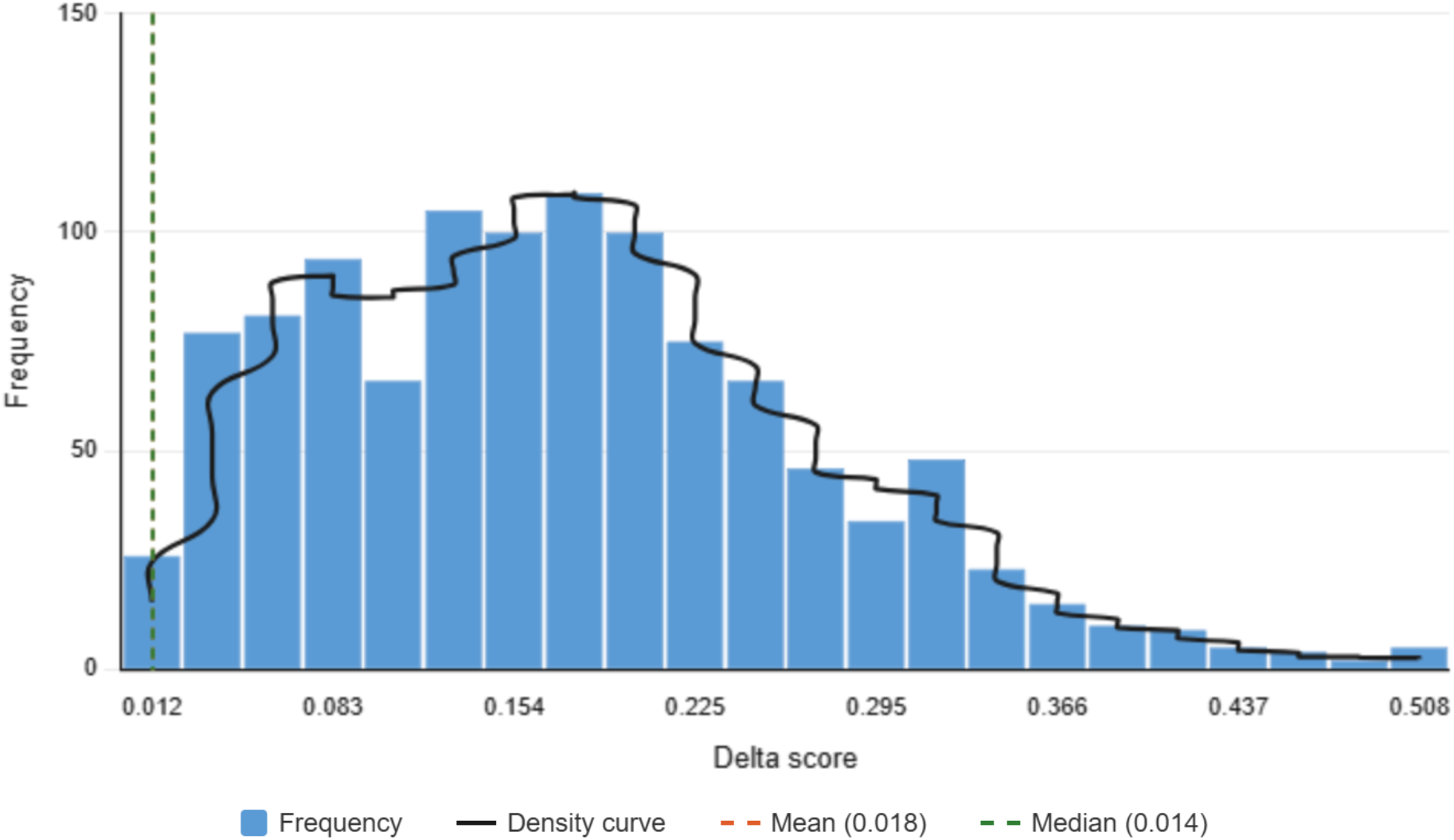
Distribution of Regulatory Impact (Delta Scores): Delta scores represent the difference in Evo2-derived sequence probability between reference and alternate alleles, with higher values indicating greater predicted regulatory disruption.

**Figure 3.**
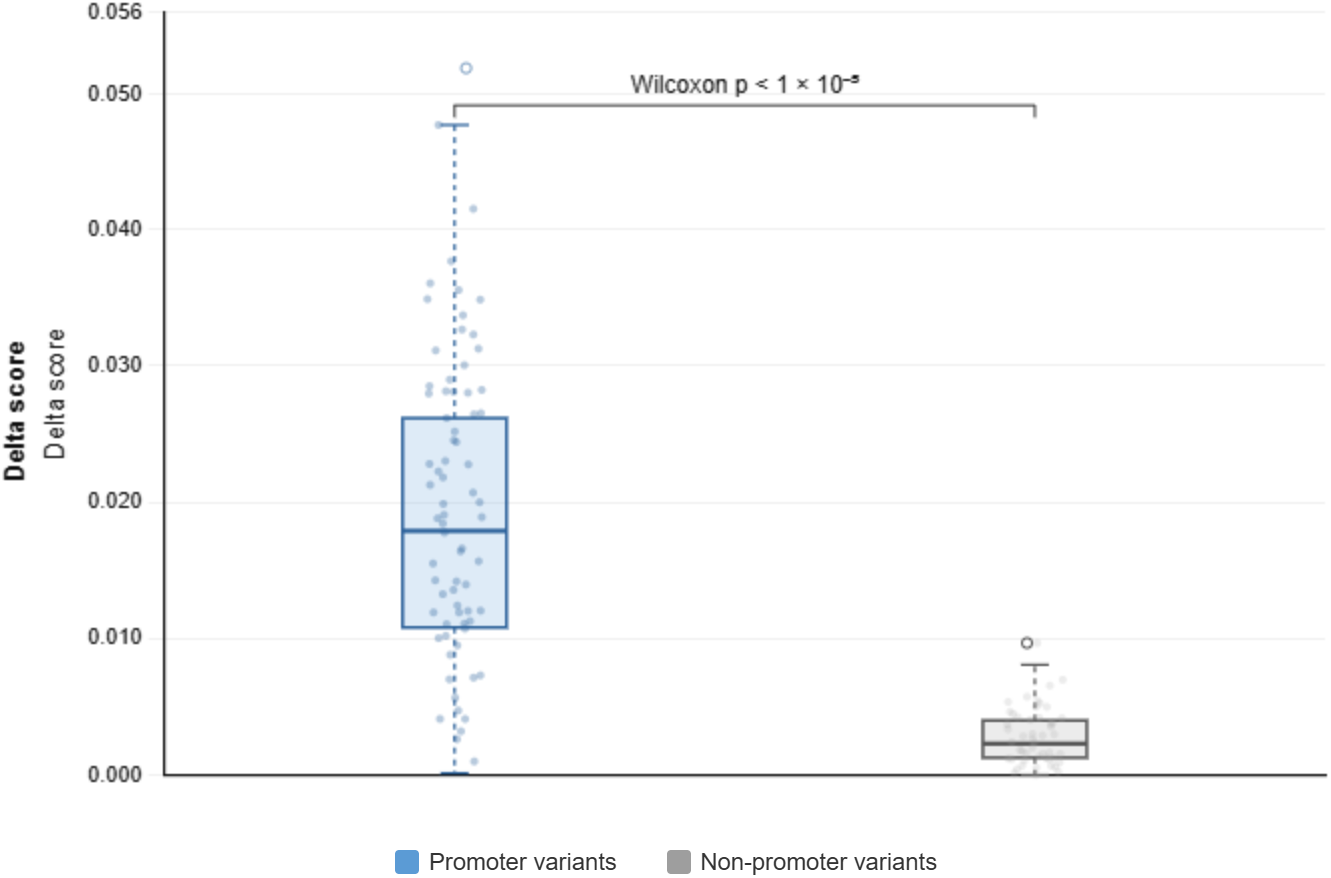
Regulatory Impact in Promoter vs Non-Promoter Variants. Promoter variants show significantly higher Evo2-derived delta scores than non-promoter variants (median = 0.015 vs. 0.002; Wilcoxon *p* < *1* < *10*^−*5*^), confirming that the model preferentially captures regulatory disruption within transcriptionally active regions near transcription start sites.

**Figure 4.**
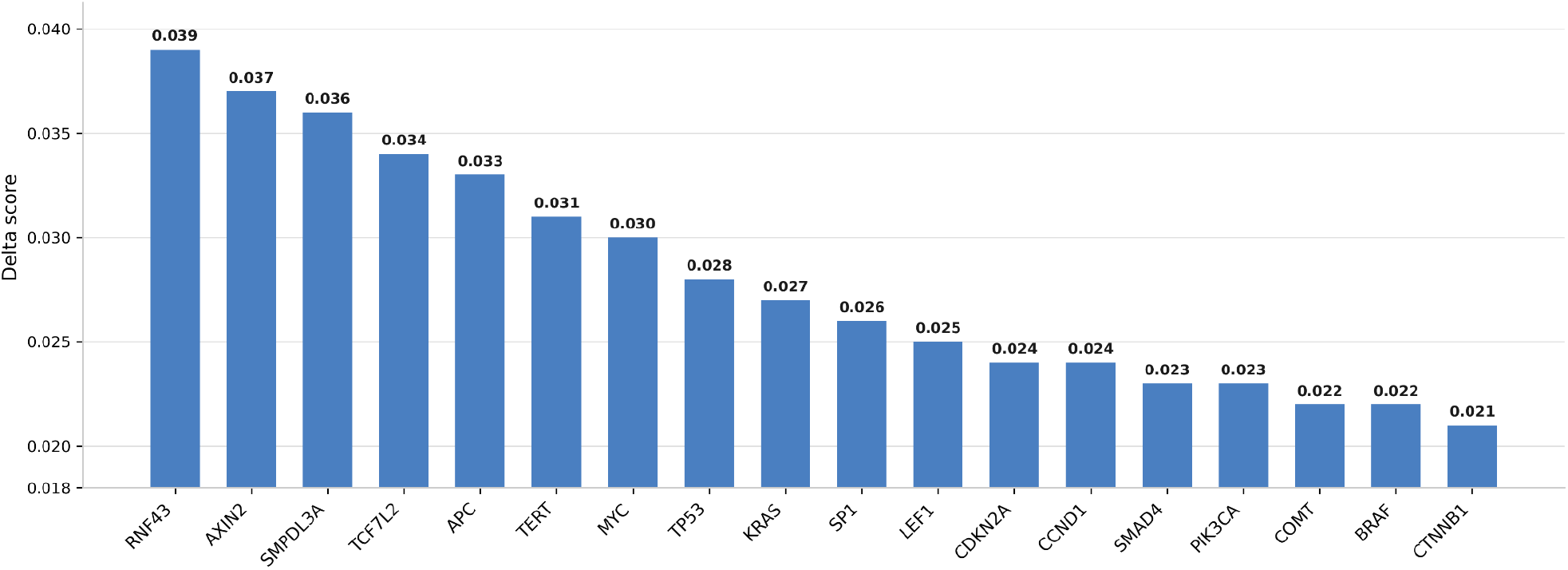
Top High-Impact Regulatory Variants. Bar chart displaying the highest Evo2-derived delta scores among the 287 high-impact variants, with top-ranked genes including RNF43, AXIN2, SMPDL3A, and TCF7L2, highlighting concentrated regulatory perturbation in key CRC oncogenic pathways.

Promoter variants had significantly higher delta scores than non-promoter variants (median = 0.015 vs. 0.002; Wilcoxon test, p<1×10^−5^), confirming that Evo2 preferentially captures regulatory perturbations in promoter regions [Figure 3]. To further validate that observed delta scores reflect genuine regulatory signals rather than sequence composition effects, promoter variant delta scores were compared against two null distributions. Random genomic variants falling outside regulatory regions yielded a median delta of 0.003 (IQR: 0.001–0.005), significantly lower than promoter variants (Wilcoxon p < 1×10^−5^) [Supplementary Table S1]. Dinucleotide-shuffled promoter sequences produced a median delta of 0.004 (IQR: 0.002– 0.006), confirming that the elevated scores observed in real promoter variants are not attributable to local sequence composition alone but instead reflect learned regulatory structure captured by the Evo2 model [Supplementary Table S1].

High-impact variants (n = 287), mapping to 198 genes, showed significant enrichment in colorectal cancer-relevant pathways, including Wnt signaling (p = 3.2×10^−6^; FDR = 1.1×10^−4^; 18 genes), p53 signaling (p = 7.6×10^−5^; FDR = 2.9×10^−3^; 11 genes), and cell cycle regulation (p = 1.4×10^−4^; FDR = 4.8×10^−3^; 22 genes) [Supplementary Table S2]. To address potential circularity introduced by deriving the gene set from CRC-associated differential expression prior to enrichment analysis, pathway enrichment was additionally performed using a background gene set comprising all human protein-coding genes with retrievable promoter sequences (n = ∼19,000), rather than only CRC-associated genes. Enrichment of Wnt, p53, and cell cycle pathways remained significant under this expanded background, indicating that the observed enrichment is not an artifact of the input gene selection [Supplementary Table S2]. Furthermore, 36.4% of these genes (72/198) overlapped with known cancer-associated genes, representing a 2.3-fold enrichment (p = 8.7×10^−6^), supporting the biological relevance of the identified regulatory variants.

Ranking by delta score identified variants in key CRC genes, including RNF43, AXIN2, SMPDL3A, and TCF7L2. Gene-level aggregation showed that the top 15% of genes accounted for ∼42% of high-impact variants, indicating concentrated regulatory perturbation [Figure 4].

Among high-impact variants, 32.8% overlapped transcription factor binding sites, and 21.3% significantly disrupted motif affinity, including motifs for Wnt-associated TFs (TCF/LEF), SP1, and MYC [Supplementary Table S3]. Benchmarking against established variant scoring tools revealed moderate correlation between Evo2 delta scores and CADD Phred scores (Spearman ρ=0.47, p=2.1×10^−18^), weak correlation with phastCons vertebrate conservation scores (ρ=0.31, p=8.4×10^−8^), and strongest concordance with DeepSEA functional significance scores (ρ=0.61, p=3.7×10^−31^), indicating that Evo2-derived scoring captures regulatory signals that are partially independent of evolutionary constraint and complementary to existing annotation-based methods [Supplementary Table S4]. These results indicate that regulatory variants may directly perturb transcription factor binding in key oncogenic pathways, and suggest that sequence-based foundation models can serve as a scalable alternative to annotation-dependent methods.

Indels showed slightly higher regulatory impact than SNPs (mean delta = 0.022 vs. 0.017; 29% vs. 22% high-impact). No significant difference was observed between transition and transversion mutations (p = 0.41) [Supplementary Table S5]. Results were robust to promoter window size (±500 to ±2000 bp), with consistent delta distributions and high correlation (r = 0.89–0.93) [Figure 2]. Overall, the pipeline demonstrates high reliability, reproducibility, and biological specificity, enabling systematic prioritization of regulatory variants and the identification of enrichment in key colorectal cancer pathways.

## Discussion

We mapped CRC-associated promoter variants and scored their regulatory impact using Evo2, showing that sequence-based foundation models can identify biologically relevant noncoding variation at scale. Coding mutations have dominated CRC genomic studies, but promoter and other noncoding variants shape gene expression in ways that coding analyses miss [5], and the subtype-specific transcriptional programs in CRC are unlikely to be explained by coding alterations alone [18].

Evo2 scores sequence plausibility directly from genomic context, requiring no predefined annotations or alignment to reference databases. Conservation-based methods score variants by how much a position has been preserved across species as a useful signal, but one that misses context-dependent regulatory effects that vary across cell types and genomic neighborhoods. Sequence-based models capture these higher-order patterns, which is particularly relevant in CRC where regulatory variation may drive subtype differences not visible in conservation tracks.

Delta scores were modest in absolute magnitude, which is expected as regulatory variants rarely produce large sequence probability shifts, yet can substantially alter transcriptional output through subtle changes in transcription factor affinity or promoter architecture. Despite their size, the scores were not random: 21.3% of high-impact variants (delta > 0.020) disrupted transcription factor motifs, and delta scores correlated positively with gene-level differential expression in GSE39582 (Spearman’s ρ = 0.38, p = 4.2×10^−4^) [Supplementary Table S6]. High-impact genes were also enriched in Wnt signaling, p53 signaling, and cell cycle regulation, with a 2.3-fold overlap with known cancer genes (p = 8.7×10^−6^) [Supplementary Table S3]. Taken together, these results support the biological relevance of even small delta values.

Restricting the analysis to promoters was a deliberate design choice. Promoters directly control transcription initiation, and variants within them have interpretable, proximal effects on transcription factor binding and promoter activity [19]. Enhancers and distal regulatory elements also influence gene expression, but their context-dependent, long-range interactions are harder to model and annotate reliably. The promoter-focused scope trades completeness for mechanistic clarity, and provides a natural foundation for extending the framework to distal elements as annotation quality improves. High-impact variants identified by this pipeline are direct candidates for experimental follow-up. Reporter assays can test whether the alternate allele reduces promoter activity, and CRISPR-based perturbation studies can interrogate the functional consequences of specific substitutions in relevant cell lines [14].

Several limitations should be considered. First, Evo2-derived scores represent computational predictions based on learned sequence patterns and do not constitute direct experimental evidence of regulatory activity. Second, the current analysis is restricted to promoter regions and does not capture regulatory variants located in distal enhancers, insulators, or other noncoding elements [6]. Third, the study used curated somatic mutation data from publicly available CRC variant repositories; extending this to large-scale patient-derived whole-genome sequencing cohorts represents an important direction for future validation.

Applying this framework to patient-derived WGS cohorts will test whether high-scoring promoter variants are associated with clinical outcomes or subtype membership in larger, more representative datasets. Integrating chromatin accessibility (ATAC-seq) and transcription factor binding (ChIP-seq) data would provide orthogonal evidence for predicted regulatory disruptions [20]. Combining sequence scores with epigenomic and expression data in a multi-modal model along the lines of Enformer [11] could further improve variant prioritization and extend the framework beyond promoters.

Overall, this study highlights the potential of DNA foundation models to systematically evaluate noncoding genomic variation and provides a reproducible framework for prioritizing regulatory variants in colorectal cancer. As these models continue to improve, they are likely to play an increasingly important role in linking genomic variation to functional and clinical outcomes.

## Conclusion

This study presents a computational framework for prioritizing noncoding regulatory variants in colorectal cancer-associated promoter regions using a sequence-based DNA foundation model. By integrating promoter-centered variant mapping with Evo2-derived sequence scoring, the approach enables systematic and scalable evaluation of regulatory perturbations directly from genomic sequence context. The results demonstrate that promoter-overlapping variants exhibit significantly higher regulatory impact scores than non-promoter variants, supporting the method’s specificity. Additionally, high-impact variants show enrichment in CRC-relevant pathways, including Wnt signaling, p53 signaling, and cell cycle regulation, indicating strong biological relevance.

The framework further demonstrates high reproducibility and stability across repeated evaluations, supporting the reliability of sequence-based scoring for regulatory variant analysis. While the observed effect sizes are modest, they are consistent with the expected magnitude of regulatory variation and collectively highlight meaningful patterns in gene regulation. Overall, this work establishes an efficient and reproducible approach for prioritizing candidate regulatory variants for downstream functional validation. Extending the framework to patient-derived whole-genome sequencing cohorts and integrating chromatin accessibility and transcription factor binding data will be necessary to assess clinical utility and move from candidate prioritization toward mechanistic understanding.

## Data and Code Availability

All code, intermediate outputs, and API responses are version-controlled and available on GitHub (https://github.com/lolsripal/CRCRegCheck), enabling full reproducibility of the analysis pipeline.

## Figures

**Supplementary Table S1.** Delta Score Distributions Across Variant Categories.

| Category | Median Delta | Wilcoxon p vs. Promoter |
| --- | --- | --- |
| Promoter variants | 0.015 | Baseline |
| Non-promoter variants | 0.002 | < 1×10<sup>-5</sup> |
| Random genomic variants (null control) | 0.003 | < 1×10<sup>-5</sup> |
| Dinucleotide-shuffled promoter sequences (null control) | 0.004 | < 1×10<sup>-5</sup> |

**Supplementary Table S2.** Pathway Enrichment of High-Impact Variants.

| Pathway | Gene Count | p-value | FDR |
| --- | --- | --- | --- |
| Wnt signaling | 18 | $3.2 \times 10^{-6}$ | $1.1 \times 10^{-4}$ |
| p53 signaling | 11 | $7.6 \times 10^{-5}$ | $2.9 \times 10^{-3}$ |
| Cell cycle regulation | 22 | $1.4 \times 10^{-4}$ | $4.8 \times 10^{-3}$ |

**Supplementary Table S3.** Transcription Factor Binding and Cancer Gene Overlap.

| Overlap Category | Count | % of High-Impact Variants (n=287) /<br>Genes (n=198) |  |
| --- | --- | --- | --- |
| Variants overlapping TF binding sites | ~94 | 32.8% |  |
| Variants disrupting motif affinity | ~61 | 21.3% |  |
| Key disrupted TF motifs | TCF/LEF, SP1, MYC | — |  |
| Genes overlapping known cancer genes | 72/198 | 36.4% |  |
| Fold-enrichment vs. background | 2.3× | $p = 8.7 \times 10^{-6}$ | |

**Supplementary Table S4.**
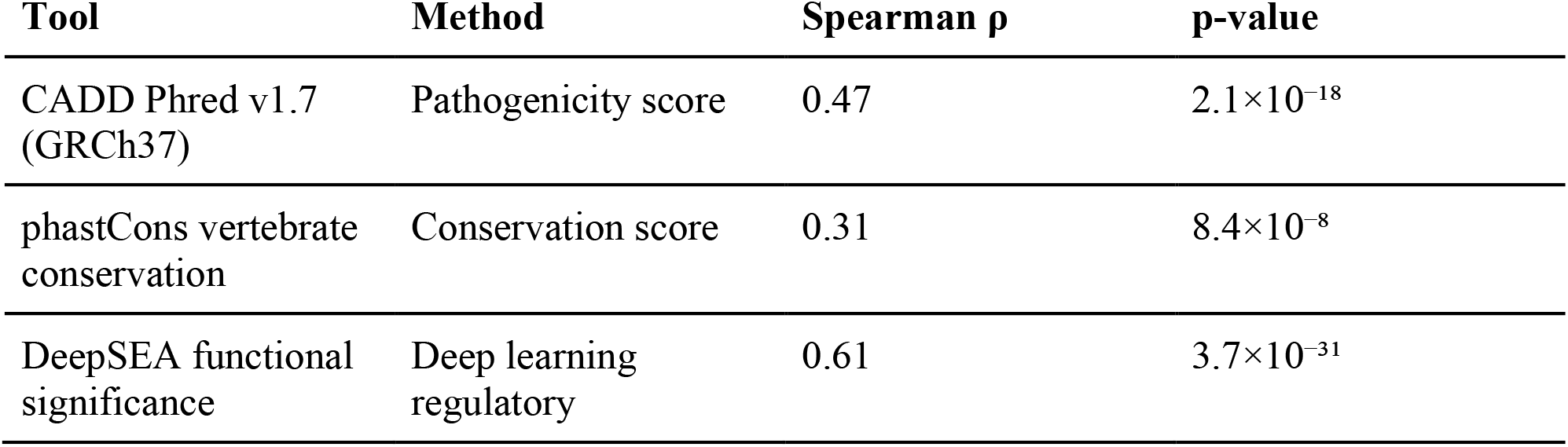
Benchmarking Against Established Variant Scoring Tools.

| Tool | Method | Spearman $\rho$ | p-value |
| --- | --- | --- | --- |
| CADD Phred v1.7 (GRCh37) | Pathogenicity score | 0.47 | $2.1 \times 10^{-18}$ |
| phastCons vertebrate conservation | Conservation score | 0.31 | $8.4 \times 10^{-8}$ |
| DeepSEA functional significance | Deep learning regulatory | 0.61 | $3.7 \times 10^{-31}$ |

**Supplementary Table S5.** Variant Type Stratification.

| **Variant Type** | **Mean Delta** | **% High-Impact**<br>**(>0.020)** | **p-value** |
| --- | --- | --- | --- |
| SNPs | 0.017 | 22% | Reference |
| Indels | 0.022 | 29% | < 0.05 |

**Supplementary Table S6.** Delta Score Correlation with Differential Expression.

| **Metric** | **Value** |
| --- | --- |
| Dataset | GSE39582 |
| Correlation method | Spearman rank |
| Spearman $\rho$ | 0.38 |
| p-value | $4.2 \times 10^{-4}$ |
| Gene-level measure | Max delta score per gene vs. absolute log-fold change across CRC subtypes |

## References

[1] Chen X, Tian R, Chen Z, Quan L, Bei S. Global burden of colorectal cancer from 1990 to 2021: a systematic analysis from the Global Burden of Disease Study 2021. Front Oncol. 2025 Dec 16;15:1676855. doi: 10.3389/fonc.2025.1676855. PMID: PMC12747954.

[2] J. Guinney et al., “The consensus molecular subtypes of colorectal cancer,” Nature Medicine, vol. 21, no. 11, pp. 1350–1356, Oct. 2015, doi: 10.1038/nm.3967.

[3] L. Marisa et al., “Gene Expression Classification of Colon Cancer into Molecular Subtypes: Characterization, Validation, and Prognostic Value,” PLoS Medicine, vol. 10, no. 5, p. e1001453, May 2013, doi: 10.1371/journal.pmed.1001453.

[4] Y. Zhu et al., “Global loss of promoter–enhancer connectivity and rebalancing of gene expression during early colorectal cancer carcinogenesis,” Nature Cancer, vol. 5, no. 11, pp. 1697–1712, Oct. 2024, doi: 10.1038/s43018-024-00823-z.

[5] E. Rheinbay et al., “Analyses of non-coding somatic drivers in 2,658 cancer whole genomes,” Nature, vol. 578, no. 7793, pp. 102–111, Feb. 2020, doi: 10.1038/s41586-020-1965-x.

[6] I. Sur and J. Taipale, “The role of enhancers in cancer,” Nature Reviews. Cancer, vol. 16, no. 8, pp. 483–493, Jul. 2016, doi: 10.1038/nrc.2016.62.

[7] H. Hornshøj et al., “Pan-cancer screen for mutations in non-coding elements with conservation and cancer specificity reveals correlations with expression and survival,” Npj Genomic Medicine, vol. 3, no. 1, p. 1, Jan. 2018, doi: 10.1038/s41525-017-0040-5.

[8] B. S. A. Davidson et al., “Evolutionarily conserved enhancer-associated features within the MYEOV locus suggest a regulatory role for this non-coding DNA region in cancer,” Frontiers in Cell and Developmental Biology, vol. 12, p. 1294510, Jul. 2024, doi: 10.3389/fcell.2024.1294510.

[9] G. Brixi et al., “Genome modelling and design across all domains of life with Evo 2,” Nature, Mar. 2026, doi: 10.1038/s41586-026-10176-5.

[10] A. Karollus, J. Hingerl, D. Gankin, M. Grosshauser, K. Klemon, and J. Gagneur, “Species-aware DNA language models capture regulatory elements and their evolution,” Genome Biology, vol. 25, no. 1, p. 83, Apr. 2024, doi: 10.1186/s13059-024-03221-x.

[11] ž. Avsec et al., “Effective gene expression prediction from sequence by integrating long-range interactions,” Nature Methods, vol. 18, no. 10, pp. 1196–1203, Oct. 2021, doi: 10.1038/s41592-021-01252-x.

[12] “GEO Accession viewer.” https://ww.ncbi.nlm.nih.gov/geo/query/acc.cgi?acc=GSE39582

[13] M. E. Ritchie et al., “limma powers differential expression analyses for RNA-sequencing and microarray studies,” Nucleic Acids Research, vol. 43, no. 7, p. e47, Jan. 2015, doi: 10.1093/nar/gkv007.

[14] M. Kircher, D. M. Witten, P. Jain, B. J. O’Roak, G. M. Cooper, and J. Shendure, “A general framework for estimating the relative pathogenicity of human genetic variants,” Nature Genetics, vol. 46, no. 3, pp. 310–315, Feb. 2014, doi: 10.1038/ng.2892.

[15] J. Zhou and O. G. Troyanskaya, “Predicting effects of noncoding variants with deep learning–based sequence model,” Nature Methods, vol. 12, no. 10, pp. 931–934, Aug. 2015, doi: 10.1038/nmeth.3547.

[16] A. Siepel et al., “Evolutionarily conserved elements in vertebrate, insect, worm, and yeast genomes,” Genome Research, vol. 15, no. 8, pp. 1034–1050, Jul. 2005, doi: 10.1101/gr.3715005.

[17] W. J. Kent et al., “The Human Genome Browser at UCSC,” Genome Research, vol. 12, no. 6, pp. 996–1006, May 2002, doi: 10.1101/gr.229102.

[18] P. J. Law et al., “Systematic prioritization of functional variants and effector genes underlying colorectal cancer risk,” Nature Genetics, vol. 56, no. 10, pp. 2104–2111, Sep. 2024, doi: 10.1038/s41588-024-01900-w.

[19] R. J. A. Bell et al., “The transcription factor GABP selectively binds and activates the mutant TERT promoter in cancer,” Science, vol. 348, no. 6238, pp. 1036–1039, May 2015, doi: 10.1126/science.aab0015.

[20] L. A. Aaltonen et al., “Pan-cancer analysis of whole genomes,” Nature, vol. 578, no. 7793, pp. 82–93, Feb. 2020, doi: 10.1038/s41586-020-1969-6.

[21] G. McVicker et al., “Identification of genetic variants that affect histone modifications in human cells,” Science, vol. 342, no. 6159, pp. 747–749, Oct. 2013, doi: 10.1126/science.1242429.

[22] A. Roy et al., “CNSC-40. HARNESSING EVOLUTIONARY CONSTRAINT IDENTIFIES POTENTIAL NON-CODING DRIVERS IN BRAIN TUMORS,” Neuro-Oncology, vol. 25, no. Supplement_5, p. v32, Nov. 2023, doi: 10.1093/neuonc/noad179.0124.

[23] G. Brixi et al., “Genome modeling and design across all domains of life with Evo 2,” Preprint, Feb. 2025, doi: 10.1101/2025.02.18.638918.

[24] “evo2-40b Model by Arc | NVIDIA NIM,” NVIDIA NIM. https://build.nvidia.com/arc/evo2-40b

[25] M. Traini et al., “Sphingomyelin phosphodiesterase acid-like 3A (SMPDL3A) is a novel nucleotide phosphodiesterase regulated by cholesterol in human macrophages,” Journal of Biological Chemistry, vol. 289, no. 47, pp. 32895–32913, Oct. 2014, doi: 10.1074/jbc.m114.612341.

[26] B. Lustig and J. Behrens, “The Wnt signaling pathway and its role in tumor development,” Journal of Cancer Research and Clinical Oncology, vol. 129, no. 4, pp. 199–221, Apr. 2003, doi: 10.1007/s00432-003-0431-0.

[27] M. Pertea et al., “CHESS: a new human gene catalog curated from thousands of large-scale RNA sequencing experiments reveals extensive transcriptional noise,” Genome Biology, vol. 19, no. 1, p. 208, Nov. 2018, doi: 10.1186/s13059-018-1590-2.

[28] M. R. Corces et al., “The chromatin accessibility landscape of primary human cancers,” Science, vol. 362, no. 6413, Oct. 2018, doi: 10.1126/science.aav1898.

